# Race to survival during antibiotic breakdown determines the minimal surviving population size

**DOI:** 10.1101/2022.08.04.502802

**Authors:** Lukas Geyrhofer, Philip Ruelens, Andrew D. Farr, Diego Pesce, J. Arjan G.M. de Visser, Naama Brenner

## Abstract

A common strategy used by bacteria to resist antibiotics is enzymatic degradation or modification. Such a collective mechanism also enhances the survival of nearby cells, an effect that increases with the number of bacteria that are present. Collective resistance is of clinical significance, yet a quantitative understanding at the population level is lacking. Here we develop a general theoretical framework of collective resistance under antibiotic degradation. Our modeling reveals that population survival crucially depends on the ratio of timescales of two processes: the rates of population death and antibiotic removal. However, it is insensitive to molecular, biological and kinetic details of the underlying processes that give rise to these timescales. Another important aspect for this ‘race to survival’ is the degree of ‘cooperativity’, which is related to the permeability of the cell wall for antibiotics and enzymes. These observations motivate a coarse-grained, phenomenological model and simple experimental assay to measure the dose-dependent minimal surviving population size. From this model, two dimensionless parameters can be estimated, representing the population’s race to survival and single-cell resistance. Our simple model may serve as reference for more complex situations, such as heterogeneous bacterial communities.

## 1 Introduction

Antibiotic resistance is an outstanding global health problem [1, 2]. Much research has been devoted to understanding the molecular mechanisms utilized by bacteria to resist antibiotics, and multiple resistance and tolerance mechanisms have been discovered and described in the past decades [3]. Antibiotic resistance poses serious problems when resistant cells spread in space and time through a population. The coupling of molecular mechanisms with population dynamics is therefore crucial for deepening our understanding of resistance and ultimately to control its spread.

It is increasingly apparent that antibiotic resistance also depends on population-level effects, which can be broadly categorized into two classes. In the first class, the resistance of one cell positively affects the survival of other cells nearby. This collective (or cooperative) resistance involves decreasing the effective concentration of antibiotics in the environment, for example via binding to cellular components of cells or enzymatic degradation [4, 5]. In the second class, efforts of an individual cell to resist antibiotics might harm its neighboring cells, for example via efflux pumps that keep the internal antibiotics concentration low at the expense of local increases of the antibiotic concentration surrounding the resistant cells.

In this work, we concentrate on the first, collective form of antibiotic resistance. In particular we focus on the production of enzymes that degrade or modify antibiotics, rendering them ineffective. In some cases these enzymes are secreted outside the cells [6, 7], while in others, most of the degradation takes place inside the cell [8]. *β*-lactamases are the best known enzymes implementing such a degradation strategy by hydrolyzing *β*-lactam antibiotics, both inside the periplasm and leaking out of the cell through outer-membrane porins [9, 7], but several other examples are also known [10, 11]. The result is a gradual removal of antibiotic in the environment, which potentially alleviates stress and aids the survival of nearby cells. This helps resistant cells to survive and establish a population [12, 13]. However, and perhaps more conspicuously, this strategy may also enhance the survival of nearby sensitive (non-resistant) cells, as once the antibiotic concentration has been reduced below their threshold for growth [8, 14], the sensitive population can expand and even compete for resources with the resistant cells [15, 16, 17].

In collective resistance mechanisms, the size of the population matters: more cells produce more degrading enzyme, which relieves antimicrobial stress more rapidly and enhances the probability of recovery of the population before its extinction. Additionally, the time-window available before extinction is directly dependent on population size. Therefore, collective resistance exhibits an inoculum effect [18] and the concept of a standard MIC - the Minimal Inhibitory Concentration for bacterial growth - is no longer well-defined, since the success of overcoming the antibiotics at a given concentration depends on cell number. Previous work has addressed this inoculum effect with several different approaches. Artemova et al. [19] studied evolutionary fitness and found that it is the single-cell MIC (scMIC) defined for an isolated cell, rather than the standard inoculum-dependent MIC, which mostly determines fitness of a resistant strain and is a better measurement for predicting the evolution of resistance. More recently several kinetic models were tested to describe measurements of inoculum-dependent MIC in different cases [20]. For example, Saebelfeld et al. [17] used a simple branching model to predict the MIC of resistant strains in the absence of social interactions, as a reference to detect collective resistance.

Here, we are interested in the population dynamics of collective antibiotic resistance, and highlight the interplay between cooperative and selfish aspects of such resistance. In particular, an enzyme that hydrolyzes or modifies antibiotics could be public if it is excreted and shared across the environment, or private if it remains intracellular and degradation happens inside the cell. In between, partial enzyme excretion constitutes an intermediate level of privatization of antibiotic resistance via enzymatic degradation. Our modeling framework allows for quantifying such a varying level of privatization in collective resistance. We describe the sensitivity of a population to antibiotics by an inoculum-dependent MIC, or alternatively, a minimal surviving population size (MSPS) that can overcome a given antibiotic concentration. Using mathematical modeling, we show that the shape of this MSPS curve has universal features and is only weakly dependent on molecular details and reaction kinetics. Rather, it reflects a relationship between the timescales of the population death rate, the antibiotic removal rate, and the level of privatization of the antibiotic-removal mechanism. We propose a simple experimental assay to determine the MSPS curve and show that our predictions are in good agreement with experimental data.

## 2 Model

### 2.1 Dynamic model for collective race-to-survival

In a bacterial population, the number of cells *N* (*t*) grows exponentially with time *t* with growth rate *α* as

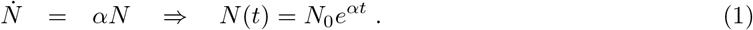

The effect of antibiotics on a population can be described by a dependence of its growth rate on antibiotics concentration *B*. Fig. 1A shows this dependence *α*(*B*) measured for *E. coli* cells exposed to different concentrations of cefotaxime (CTX), a cephalosporin class *β*-lactam. In the absence of antibiotics (*B* = 0), the population growth rate *α*_0_ reflects the specific bacterial strain and the nature of environment, e.g. medium composition. Increasing antibiotics concentration reduces this growth rate sharply around a threshold concentration *B* ≈ *µ*. Above this threshold, growth rate *α* becomes negative as cells are killed by the antibiotics and the population size decreases. Increasing further the antibiotic concentration, it has been observed that the rate of death levels out to a constant rate proportional to the growth rate in the absence of antibiotics [21, 22].

**Figure 1:**
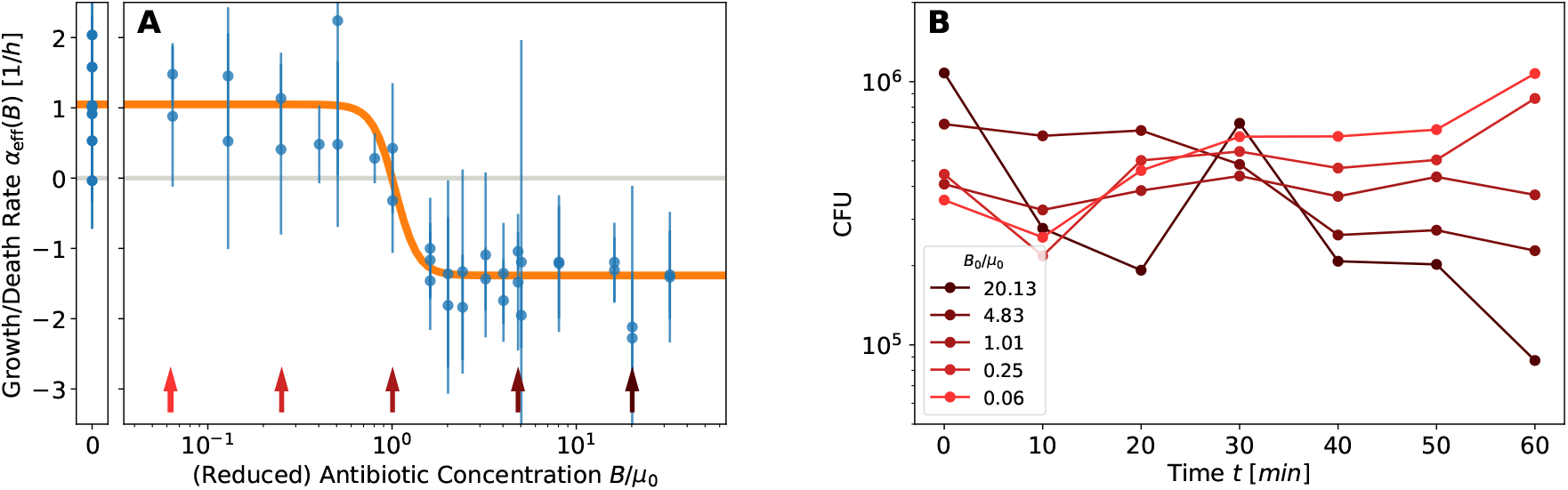
Effect of cefotaxime (CTX) on growth of sensitive *E. coli* populations using conventional kill curves. (A) Growth rate *α*(*B*) is plotted as a function of scaled CTX concentration *B* in the medium. In the absence of CTX, (*B* = 0; left narrow panel), bacteria grow with a rate *α*_0_ which depends on environmental conditions. Around the threshold of *B* ≈ *µ*, growth rate decreases sharply, and then saturates at a negative rate −*γα*_0_. Each data point in Fig. 1A is extracted from a CFU count as a function of time, a few examples of which are shown in Fig. 1B. (B) Averaged trajectories of CFU counts at selected antibiotic concentrations are shown, colored correspondingly to the arrows in the left panel.

It has been found in experiments that a good mathematical description of this dependence is a decreasing sigmoidal curve (a Hill function) [23],

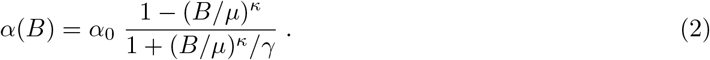

Here *µ* is the concentration at which growth decreases to zero and turns into death; *κ* the steepness of the decrease around this threshold; and −*γα*_0_ the maximal death rate. This formulation helps to separate the dependence of growth on medium or strain, through *α*_0_, from the pharmacodynamics of the antibiotics described by the sigmoidal function.

Consider a finite population placed in a closed environment with growth medium and an initial concentration of antibiotics *B*_0_, larger than the threshold *µ*. As cells are killed and the population size starts to decrease, the remaining cells produce enzymes that degrade or deactivate antibiotics. These two processes define a dynamic race-to-survival in which antibiotics must be reduced below threshold before all cells have been killed. The success of this strategy depends on the relative rates of these two processes, but also on the initial population size and initial amount of antibiotics. A larger initial population extends the time-window to achieve this goal, while more antibiotics decrease the time for the population to fight back before it goes extinct.

The reaction that reduces antibiotic concentration *B* is implemented in different strains and conditions by various kinetic processes. To illustrate the race to survival, we consider the production of an antibiotic-degrading enzyme *E* with a rate of enzyme production *ρ* proportional to the bacterial population size *N*. In turn, this enzyme degrades antibiotics *B* in a first-order biochemical reaction with catalytic efficiency *ϵ*:

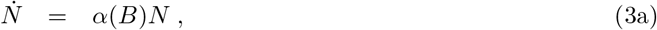

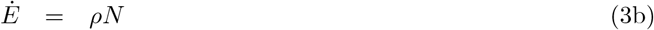

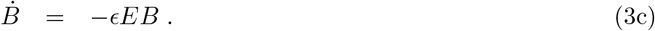

With the model of Eqs. (3), we can integrate *N* (*t*), *B*(*t*) and *E*(*t*) over time. Examples of numerically obtained trajectories are depicted in Fig. 2 for *N* (*t*), with various initial antibiotic concentrations (colors) and inoculum sizes (panels). The most crucial property determining the population’s fate is whether or not it drops below a single cell, *N* (*t*) < 1, indicating its extinction. This is the lower limit shown in the figure; if any trajectory decreases below this limit, the population goes extinct. It can be seen that for small antibiotic concentrations (orange and light brown curves) the population increases exponentially and will continue to do so until it reaches saturation (not modelled here). In contrast, for larger antibiotic concentrations (darker purple to blue curves) it eventually drops below one cell and becomes extinct. Interestingly, for larger inoculum size, at intermediate antibiotics concentration the population starts to decrease, but as antibiotic is degraded then turns around and succeeds to grow. Examples of this behavior can be found in Fig. 2.

**Figure 2:**
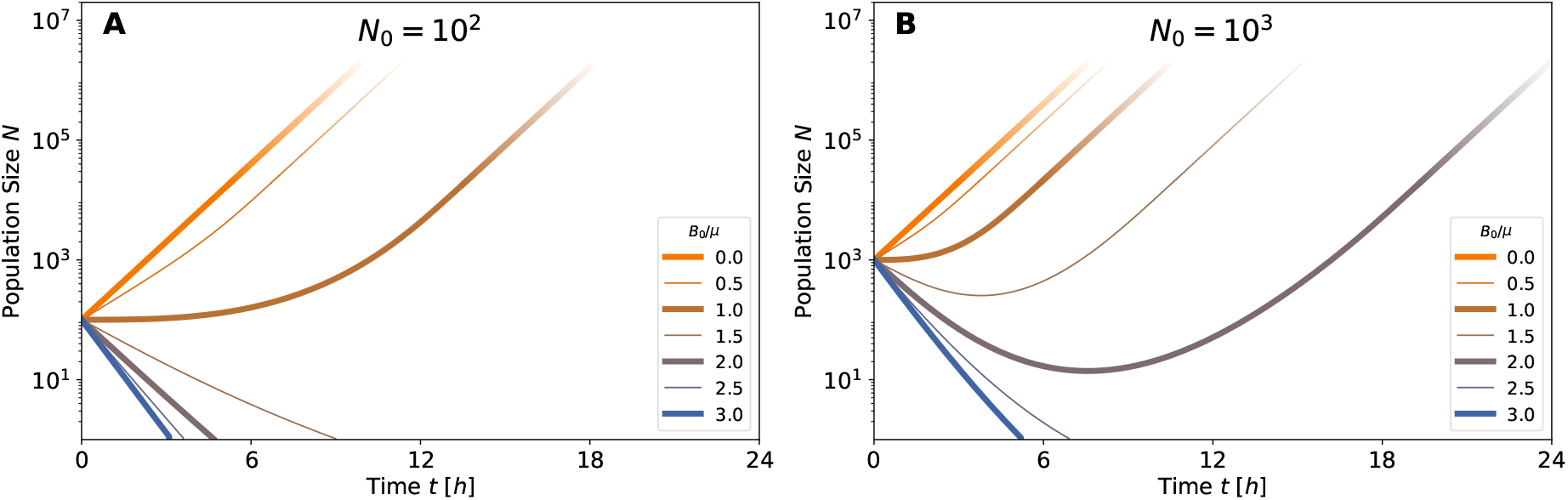
Survival depends on inoculum size and antibiotics concentration. Population size across time, *N* (*t*), computed numerically from the model (3), with various parameters: inoculum size of *N*_0_ = 10 (A), 10 (B), 1000 (C); relative initial antibiotic concentration *B*_0_*/µ* between 0 and 3 (line colors; see legend). For *B*_0_*/µ >* 1 the bacterial population initially decays, but may recover depending on inoculum size. For example, at antibiotics concentration of *B*_0_*/µ* = 2 (purple line in all panels), the larger inoculum (C) survives while the smaller ones (A,B) do not. Extinction is prevented if enough enzyme is produced during the initial population decay, such that antibiotics is reduced below the threshold concentration *µ*. In all simulations *ρϵ* = 10^−3^.

### 2.2 Minimal Surviving Population Size

Our model indicates that the inoculum effect of collective resistance is dynamic and is determined by the relative contribution of two processes: cell death and antibiotic degradation. The standard measure of Minimal Inhibitory Concentration (MIC) is used to characterize a threshold of initial antibiotic concentration *B*_0_ beyond which there is no growth at long times. However, the race to survival discussed above implies that MIC depends on inoculum size [18, 20]. Turning this dependence around, it is convenient to think about the minimal surviving population size as a function of antibiotic concentration: this Minimal Surviving Population Size (MSPS) is a curve *N*_0_(*B*_0_) rather than a single quantity such as MIC.

Using the model in Eqs. (3) we can develop an approximation to estimate this MSPS curve. If the initial antibiotic concentration is large enough, most of the race-to-survival dynamics takes place when antibiotic concentration changes only slowly. Thus, we approximate the sigmoid function for large antibiotic concentrations *B* ≫ *µ*, reducing the growth dependence on antibiotics to a simple constant:

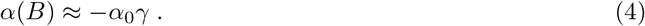

The approximation in Eq. (4) allows to find solutions to our model, which in turn provide a expression of the MSPS curve, *N*_0_(*B*_0_) (see Appendix A for derivation):

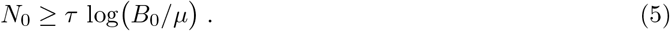

The MSPS curve, as shown in Fig. 3 (red lines), is an increasing function: when the initial antibiotic concentration increases, a larger inoculum is needed to ensure survival. Its simple form is largely robust with respect to the details of the mechanism for antibiotic degradation or inactivation (see Appendix A).

**Figure 3:**
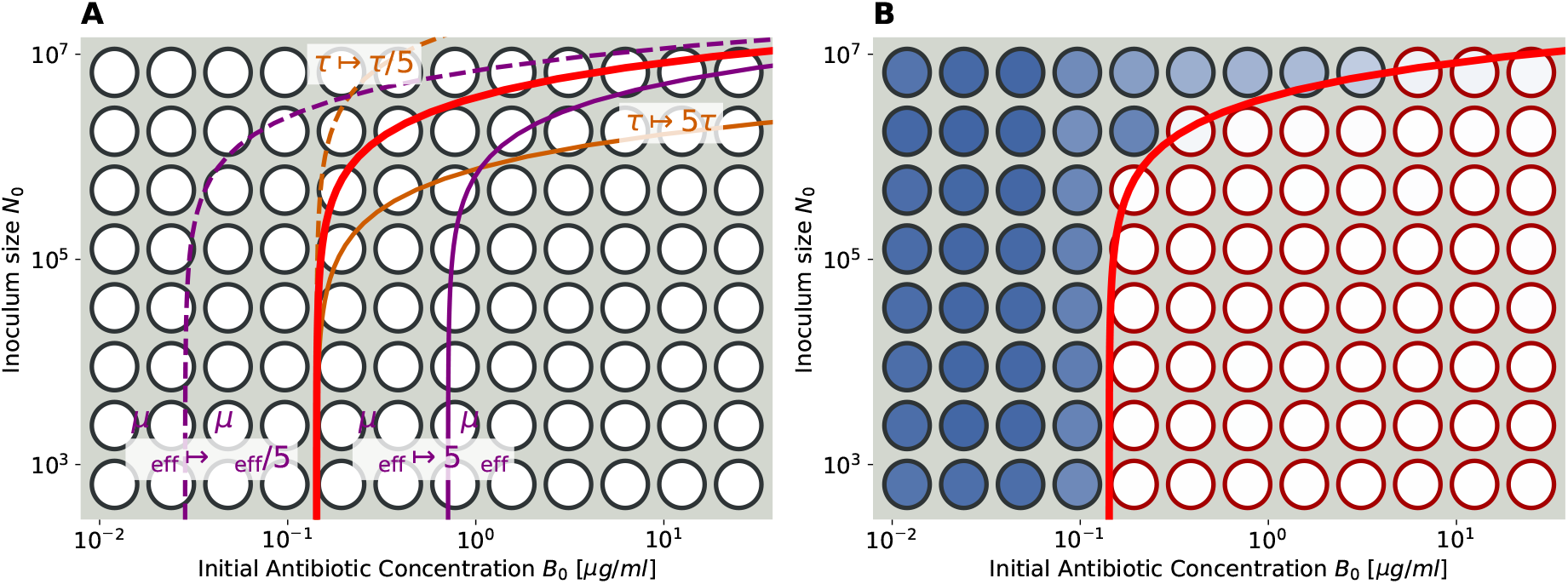
The universal MSPS curve: parameters and experiment. (A) The shape of the MSPS curve, Eq. (5), plotted in logarithmic axes (red curve). This universal curve only depends on its two parameters *τ* and *µ*_eff_. Increasing *τ* (brown curves) stretches or compresses the MSPS curve in the direction of inoculum size (y-axis). Changes in the parameter *µ*_eff_ (purple curves) induce a shift of the MSPS curve in the direction of antibiotic concentration (x-axis). (B) Experimental measurement of the MSPS curve with data from a G238S mutant: A microwell plate is started with serial dilutions of the inoculum size *N*_0_ and antibiotic concentrations *B*_0_ along the two axes. The threshold for survival is apparent by color after a long time of growth: blue wells have a population in them, while white cells do not. If the ranges of dilutions are chosen appropriately, the MSPS curve (red line) appears with its typical universal shape as the boundary between wells with surviving an extinct populations.

Let us consider the interpretation of the two parameters determining the MSPS. In our theory, *µ* is the threshold antibiotic concentration allowing growth (see Fig. 1), which sets the scale for antibiotic concentration. In terms of the plots presented in Fig. 3, *µ* is the intercept of the MSPS curve at the antibiotic concentration axis, corresponding to the small-inoculum limit (single-cell MIC; see [19]).

In addition, the MSPS depends on the dimensional parameter

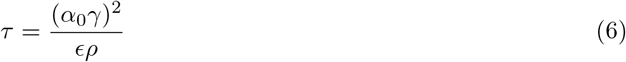

representing the ratio of timescales between killing dynamics, (*α*_0_*γ*)^2^, and the rates involved in antibiotics degradation, *ϵρ*. Large *τ* corresponds to fast killing of bacteria relative to degradation, and thus results in a higher MSPS. In terms of the plot of Fig. 3, the parameter *τ* shifts the curve along the population size axis, see Fig. 3A.

If the initial antibiotic concentration smaller, we can employ a different approximation for the growth rate. In appendix A.2 we show that 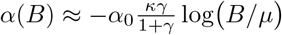, which is valid close to *B* ≈ *µ*, still allows to find solution to our model. However, a correction close to *B* ≈ *µ* should not affect the shape of the MSPS curve too much, as the asymptotic behaviour for large *B*_0_ is still given by Eq. (5).

Importantly, although we developed Eq. (5) for a specific model of enzyme degradation, the shape of the MSPS curve turns out to be insensitive to many details of the kinetics of collective resistance, as we show in Appendix A. In several example mechanisms we have analyzed, the extant parameter *τ* depends on combinations of underlying molecular parameters for different kinetic mechanisms. Nevertheless, its interpretation is always the same: *τ* represents the ratio between the rate of population death *α*_0_*λ* and the rate of reduction in antibiotic concentration.

### 2.3 Single cell privatization of resistance

Until now we have assumed that resistance is completely cooperative - produced enzymes are secreted to the common environment and directly affect the global antibiotic concentration. In this view, the antibiotic concentration is homogeneous across space and hence the same for the cell which produces the resistant enzymes and any other nearby cells which benefit from this production. In reality, at least part of the degradation takes place inside the cell (often in the periplasmic space), thus providing an increased benefit to the producing cell relative to its neighbors. In environments which are not mixed rapidly enough, even if enzymes are excreted they are more abundant in the vicinity of producing cells and will take time to diffuse away, again providing an increased benefit to the secreting cell. Our goal is now to quantify the level of privatization (or inversely, of cooperation) in collective antibiotic resistance. A full model should include transport and diffusion of concentrations in space; as a first approximation, we include the distinction between internal and external concentrations of both enzyme and antibiotics: *E*_in_, *E*_out_, *B*_in_, *B*_out_ (Fig. 4). The coupled kinetic equations for these variables can be found in Appendix B. We assume that concentrations inside the cell equilibrate much faster than outside, corresponding to its volume being tiny relative compared to the seemingly infinite reservoir of the external environment. This assumption allows to estimate the stationary internal concentrations as

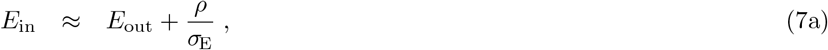

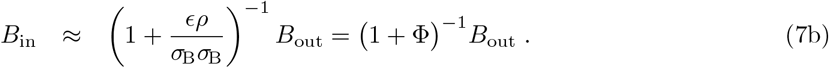

**Figure 4:**
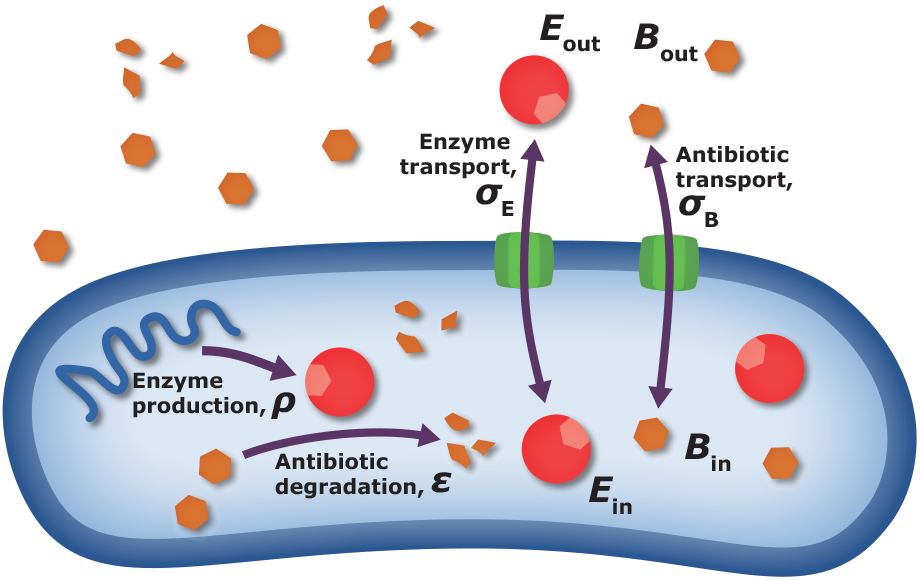
Factors affecting privatization of resistance. To describe different levels of privatization/cooperativity in antibiotic resistance, we include in our model external concentrations of both enzyme and antibiotics. The transport between external and internal spaces is determined by transport coefficients *σ*_E_, *σ*_B_. Enzyme production and antibiotic degradation efficiency are determined by *ρ, ϵ* respectively. The level of privatization is determined by the relative importance and speed of these processes.

Here, *σ*_E_, *σ*_B_ are the (linear) transport coefficients for enzyme and antibiotics respectively, passing through the cell wall. High transport coefficients will work against maintaining a concentration difference across the cell wall, and thus will promote cooperativity, while low transport coefficient will promote privatization. The effect of transport on enzyme concentration is seen in Eq. (7a), which describes the difference between internal and external enzyme concentration: internal is always larger, and high production *ρ* or alternatively low transport *σ*_E_ work to increase this difference. We can also assume that external enzyme concentrations are usually much smaller than internal concentrations.

Antibiotic concentration, in contrast, is always lower inside the cell, by a dimensionless factor that depends on both production and transport parameters. Once again, high production and low transport support a large difference, as is reflected in Eq. (7b). It is convenient to define the compound parameter appearing in Eq. (7b) as a “privatization parameter” Φ = *ϵρ/*(*σ*_B_*σ*_E_). High privatization occurs for Φ ≫ 1, corresponding to very high enzyme production *ρ*, high catalytic efficiency *c*, but low transport coefficients. In this regime, cells combat the antibiotics individually, primarily by lowering their internal concentration and not sharing degrading enzymes with neighbors. At the other end, Φ ≪ 1, the concentration is almost equal inside and outside the cell and the battle occurs in the public domain of the shared environment, resulting in maximal cooperativity to all cells and we effectively return to the first, naive model, where no distinction was made between internal and external concentrations. The privatization parameter Φ enables to interpolate between these two extremes. As usual in our modeling approach, it is not a mechanistic parameter but an effective one; ultimately, the level of privatization is determined by the difference in concentration, resulting from enzyme efficiency, production, transport, or possibly other microscopic processes.

In a real experimental setting, it is difficult – or even impossible – to measure internal concentrations of drug and enzyme, and usually only external concentrations are measurable. Nevertheless, it is the internal concentration which directly affects how antibiotics alter growth of a cell or cause its death. As can be seen from Eq. (2), the effect of antibiotics on growth rate is measured in units of *µ* and always appears as *B/µ* in all instances. Thus, Eq. (7b) allows us to translate between these two concentrations, and to write the effective growth rate as a function of the observable external antibiotic concentration:

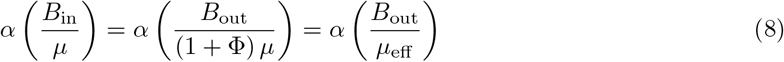

The discrepancy between (unmeasureable) internal and (measurable) external concentrations can be attributed to a modification of the growth threshold *µ*, which we define as

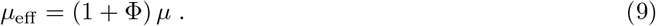

With this result, we need to modify the MSPS curve obtained in the previous section, which can now be stated in terms of the quantity *B*_0_ = *B*_out_(*t* = 0):

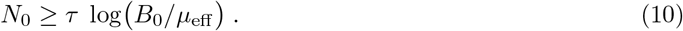

This MSPS curve in Eq. (10) has the same functional form as before with two fitting parameters *τ* and *µ*_eff_, but now one of them also includes transport properties; it is an effective parameter that deals with the difference between external and internal concentrations.

The structure of the privatization parameter Φ allows to develop our theory further. We have encountered *τ* as the ratio between the timescales of cell death (*α*_0_*λ*) and antibiotic degradation (that includes enzyme expression *ρ* and kinetic efficiency *ϵ*). The transport processes for both antibiotics and enzyme generate a third timescale in our model. Thus, we may write the privatization parameter as

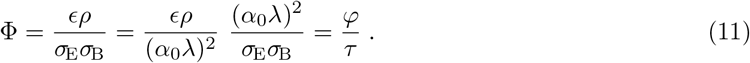

This representation is useful, because it separates intrinsic bacterial properties represented by *φ* from parameters of the antibiotic degradation mechanism. Using the expression, Eq. (11), predicts a relation between the two fitting parameters of the MSPS, *τ* and *µ*_eff_ :

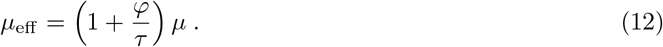

While each experiment will result in a MSPS curve with different parameters, they are not expected to be randomly scattered in the plane (*µ*_eff_, *τ*), but rather to be located on a curve defined by the Eq. (12).

## 3 Experimental validations

An experimental procedure that measures the MSPS curve is conceptually straightforward to do with a 96-well assay. Populations are inoculated with increasing inoculum size *N*_0_ along one axis, and with increasing antibiotic concentrations *B*_0_ along the other (Fig. 3B). Concentrations along the axes are chosen appropriately, such that the shape of the MSPS curve fits on the plate (which requires some prior knowledge about the ranges where the MSPS curve lies). Following overnight incubation of the cultures, a clear difference is visible between surviving and non-surviving populations. As an example, Fig. 3B shows all OD measurements of wells in different shades of blue (white indicating no growth), where an automatic threshold detection distinguishes between survival (dark gray borders) and extinction (red borders). Subsequently, one may fit Eq. (10) with its two parameters, *τ* and *µ*_eff_, at this transition from surviving to non-surviving populations on the plate. In Fig. 3 it is depicted by a red line. All steps in our algorithm to extract parameters from OD measurements of plates are described in Appendix D.2. The first parameter, *τ*, multiplies the MSPS curve and mostly determines the intersection on the *N*_0_ axis of the plate for large antibiotic concentrations. The second parameter, *µ*_eff_, corresponds to the intersection of the curve on the *B*_0_ axis for small initial population sizes, i.e. the single-cell MIC [19].

We applied this experimental procedure to several *E. coli* strains expressing various *β*-lactamases. In a first set of experiments, we used mutants of *E. coli* MG1655 with a chromosomally integrated gene encoding and expressing *β*-lactamase TEM-1 at a single intermediate level. These include a consecutive series of mutations in the TEM-1 enzyme (wildtype, G238S, G238S-E104K, G238S-E104K-M182T), which exhibit increasing catalytic efficiency *ϵ* for cefotaxime [24]. A second set of experiments used *E. coli* BW27783, where the TEM-1 gene is located on plasmid pBAD322T and its expression is controllable using an arabinose-inducible promotor. This allows to manipulate enzyme production *ρ*; the subset of mutations in TEM-1 again alters the catalytic efficiency *c* to moderate and high values. A detailed experimental protocol is described in Appendix D.1.

In this experimental setup only the gene (including its promotor) for TEM-1 is changed within both sets of experiments. Thus, we can expect that the transport properties, which have been separated in Eq. (12) as parameter *φ* should be unchanged. Ultimately, this is the reasoning behind deriving the expression Φ = *φ/τ*.

For most experiments, the MSPS curve can be fit nicely to the data and the effective parameter estimates inferred. An example of such a fit is shown in Fig. 3B. In some cases, the low resolution of a 96-well plate caused a large uncertainty in parameter values. Nevertheless, our repeated experiments yielded similar parameter values in practically all tested cases.

The resulting fitting parameters, as estimated for our collection of experiments, are shown in Fig. 5. The scaling relation predicted by the model, Eq. (12), is depicted as a gray line. Panel A shows the results of our first set of experiments with TEM-1 and its three mutants at a single expression level. The data points are not scattered in the plane, and follow the predicted curve to a good extent with TEM-1, the single, double and triple mutant being positioned increasingly higher along the curve mirroring their respective increase in resistance level [24]. Panel B presents the results for the second set of experiments exhibiting different expression levels of two TEM variants (single and triple mutant). While the single mutant follows our prediction, the triple mutant deviates from the curve at medium and high expression levels. We speculate that this deviation could arise due to heavy stress on the bacteria from the over-expression of *β*-lactamase, which significantly lowers their overall growth rate *α*_0_ (data not shown). Moreover, it may be that the external enzyme concentration is not negligible and the approximation Eq. (7b) is invalid. In these cases the relation between the two fitting parameters is not expected to obey the scaling relation. However, in general, the set of experiments clearly do not scatter randomly in the parameter plane, but are strongly correlated with one another, and – except for the two outlying points – are well described by our model.

**Figure 5:**
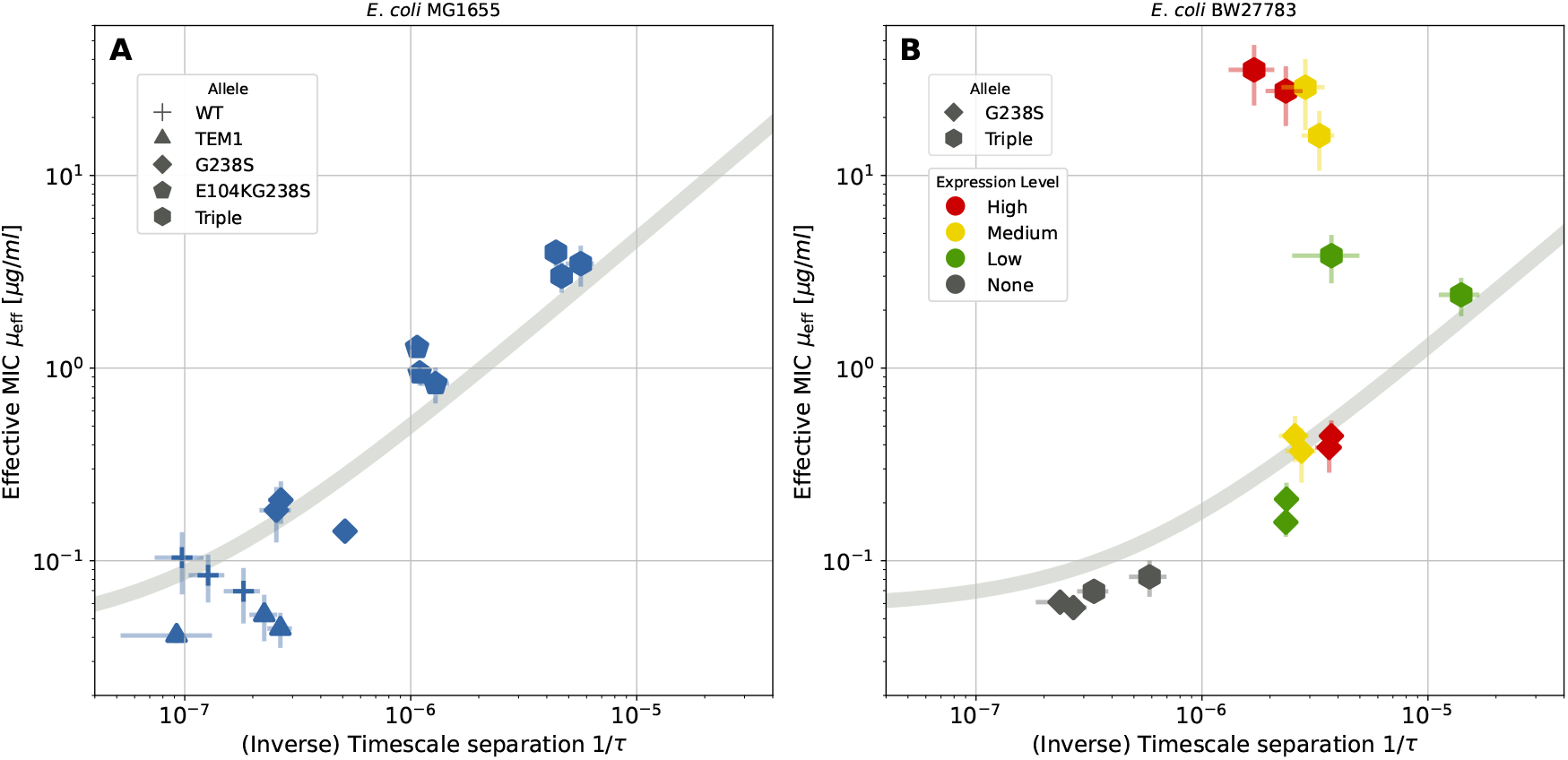
Correspondence between *µ*_eff_ and compound parameters inferred from measured MSPS curves using *β*-lactamase producing strains with varying resistance levels. (A) Best fit of Eq. (9)) for strains expressing TEM-1 variants with different catalytic efficiency *ϵ* towards cefotaxime. (B) Best fit of Eq. (9) for a strain with low (G238S, rhombus) or high catalytic efficiency *ϵ* (Triple, hexagon) and different expression levels *ρ*. Error bars are estimated in the fitting algorithm (see Appendix D.2.2).

## 4 Discussion

Collective resistance is a phenomenon by which a bacterial population can survive an antibiotic dose which a single bacterium can not [16]. This collective behaviour may have a profound influence on the effectiveness of an antibiotic therapy and thus may pose serious health risks. Using mathematical modelling, we studied collective resistance via the common resistance mechanism of antibiotic degradation or modification, and proposed a unifying framework to describe the minimal population size needed to survive a specific antibiotic concentration. We developed a conveniently simple approximation of our model that allows us to determine the dose-dependent minimal surviving population size (MSPS) curve. This curve is a generally increasing function, with a larger inoculum being able to survive increasingly higher antibiotic concentrations. Interestingly, we showed that basic features of this curve, and by extension of collective resistance via antibiotic degradation or modification, are insensitive to details of the exact resistance mechanism. Rather than kinetic parameters, the central quantities determining survival are ratios between timescales, when populations race against time to survive by collectively decreasing the antibiotic concentration.

A first parameter central to the MSPS curve describes the ratio of timescales between population extinction by antibiotic killing and antibiotic degradation. This parameter, referred to as *τ* in our model, is dimensionless and is agnostic to the exact mechanism by which the antibiotic is degraded or modified (Appendix A). This is in line with previous work [25], which also reported that it is hard or even impossible to determine details of the degradation mechanism by only observing microbial population dynamics. The hiding of microscopic kinetic details from higher population level dynamics is a form of buffering between levels of organization. While insights into the molecular mechanisms need to be known for a targeted antimicrobial therapy to be effective, the relative robustness of population-level dynamics, as shown by our modelling, is important for a basic understanding of resistance at the population or community level. The second central feature in our model is the level of privatization of the resistance mechanism, namely how much of the degradation or modification of antibiotics takes place in the shared environment, relative to the intracellular environment. Previous work has considered the limit of maximally private degradation which takes place inside the periplasmic space [26]. Their relation between internal and external threshold concentration is similar to ours in that limit, consisting of the ratio between hydrolysis rate and transport rate. Other works incorporated directly one form of collective resistance, namely, the lysis of cells as they die and the release of their enzyme content [19]. Our general result Eq. (7b) interpolates between the high privatization limit, and the other limit of low privatization where hydrolysis takes place primarily in the public domain. This provides a coarse grained phenomenological description that could apply specifically to lysis or secretion.

Although we here studied the MSPS curve of single strains, quantifying the level of privatization may have far reaching consequences for cross-protection between microbial communities and the eco-evolutionary dynamics of antibiotic resistance [13, 27]. The relationship between single-cell and population-level MIC was characterized also in previous work, where it was found that the single-cell properties affect long-term evolutionary outcomes [19]. Nevertheless, intermediate timescale dynamics of mixed populations can also be strongly affected by the collective dynamics, highlighting the importance of the MSPS concept.

Based on our model, we proposed a simple 96-well microtitre-plate assay that allows us to characterize the parameters describing the MSPS curve. We provide a detailed protocol in Appendix D to perform this assay and extract the relevant parameters, *τ* and *µ*_eff_. This method relies on the idea that a 96-well plate can serve to scan inoculum sizes and antibiotic concentrations in parallel and provide a platform for mapping the MSPS curve. A similar idea was proposed for characterizing antibiotic persistence, where the exposure time is a critical variable and is varied along one axis of a 96-well plate by changing the time at which the medium is inoculated [28].

The assay was used to assess the model at two levels. First, we tested the fit of our predicted MSPS curve to describe the border between surviving and non-surviving populations and found good agreement. Second, we performed two sets of experiments, where we independently varied the catalytic efficiency or expression level of an antibiotic-degrading enzyme in bacterial strains. Under the approximation of significant privatization and independence of other physiological properties of the bacteria, we derived a relation between fitting parameters across the set of experiments, Eq. (12). The results show, that this relation agrees rather well for all but two data points, specifically strains expressing a *β*-lactamase with very high catalytic efficiency (the triple mutant) at high levels (Fig. 5). The current experiments do not allow us to identify the source of this qualitative discrepancy, but since the triple mutant MSPS curve was assayed at significantly higher antibiotic concentrations, this may affect cell physiology via the induction of the SOS response and *β*-lactam-induced filament formation [29], or by a varying level of lysis and release of enzyme. This would modify the privatization parameter, which the model assumes to be fixed along the curve. Moreover, differences in cost of expression between *β*-lactamase alleles could also lead to differential effects of expression on cell physiology and permeability potentially affecting the timescale ratios describing the MSPS curve.

In summary, our work contributes to identifying generic mechanism-independent parameters that can be inferred from population data. It identifies robust parameter combinations that govern population dynamics in collective resistance. Specifically, it reveals relative timescales in the race for survival of populations which inactivate antibiotics that kill them, as well as levels of cooperation vs. privatization of resources in the fight against antibiotics. Our experimental framework adds a dimension to the characterization of antibiotic resistance by a concentration threshold: it extends this notion to an inoculum-dependent threshold relevant for cells utilizing a collective resistance mechanism. Our framework is expected to be amenable for extension also to the interaction between resistant and sensitive strains, which has been partially addressed from a different perspective [15].

## A Universality in antibiotic degradation dynamics

An intriguing theoretical observation is that the MSPS curve of Eq. (5) is insensitive to the specific antibiotic decay mechanisms – it is universal to a good approximation. Below, we show that multiple degradation dynamics lead to similar expressions for the curve, utilizing only the two extant parameters *τ* and *µ*, that characterize the time scale ratio between cell death and degradation, and antibiotic concentration at zero-growth, respectively. In the main text, we only considered antibiotic degradation by an excreted extracellular enzyme. Other (bio)chemical or biological processes could provide resistance against the action of the antibiotic by reducing its concentration. A few examples of such mechanisms, stated as simple reactions (with rates indicated above arrows), are as follows,

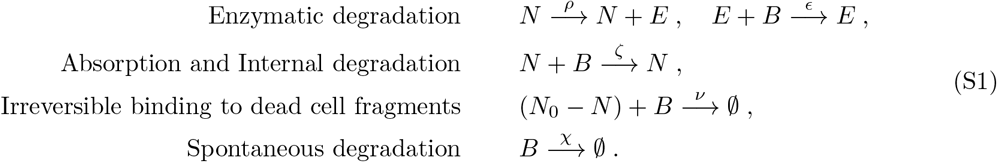

The first reaction in this list corresponds to the dynamics with excreted extracellular enzyme, with linear kinetics, discussed in the main text. In the second reaction, antibiotics is absorbed into the cell, and is degraded by some unspecified cellular reaction inside the cell. In the third example, we consider irreversible binding of antibiotics to fragments of dead cells. Their concentration is proportional to the number of cells that have died during exposure; this expression is relevant as long as population size is decreasing. Finally, we also treat spontaneous environmental degradation of antibiotics, which cells do not actively contribute to.

For all these mechanisms, the dynamics of cells and antibiotics can be described by the dynamical system,

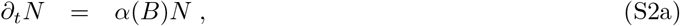

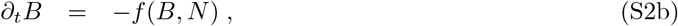

where *f* (*B, N*) specifies one of the degradation reactions of antibiotics, as listed above in Eq. (S1). Throughout this appendix, we also use the notation *∂*_*t*_ for derivations with respect to time *t*, as it helps with the ensuing calculations. Turning the reactions in Eqs. (S1) into equations, we find the different mechanisms follow the kinetics:

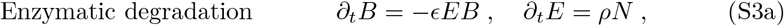

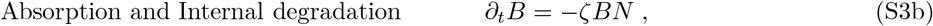

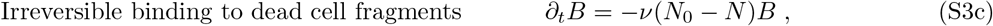

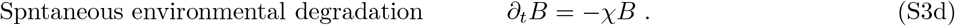

Below, we will solve these dynamical systems in detail, leading to a condition on the minimal inoculum size *N*_0_ that can recover from an initial antibiotic concentration *B*_0_:

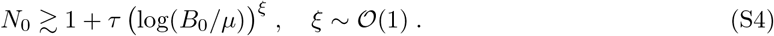

This is a general expression common to all kinetic mechanisms; it defines the universal MSPS. Each degradation mechanisms is characterized by a timescale separation *τ*, which is the ratio between two characteristic rates in that specific mechanism: a population kill rate and an antibiotic degradation rate. The exponent *ξ* varies slightly depending on the mechanism, but always assumes values close to 1. Moreover, the different approximations for the growth rate *α*(*B*) explained in the next paragraph also influence the value of *ξ*.

**Figure S1:**
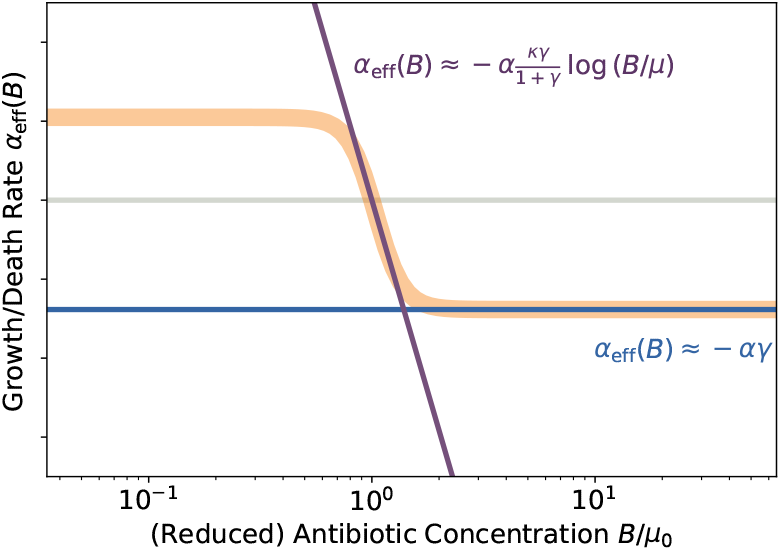
Approximations for effective growth rate. Two regions are observed, a log-linear regime around MIC and a constant death rate at antibiotic concentrations much larger that MIC.

In general, the sigmoidal function for growth rate *α*(*B*), Eq. (2), can not be used in its full form to compute solutions to the dynamics. As observed in Fig. 1, the effective kill-rate is constant for very large antibiotic concentrations. In contrast, for antibiotic concentrations close to *B/µ* ≈ 1, it is approximately linear in log(*B/µ*). Mathematically, we will deal with two approximations corresponding to these regimes:

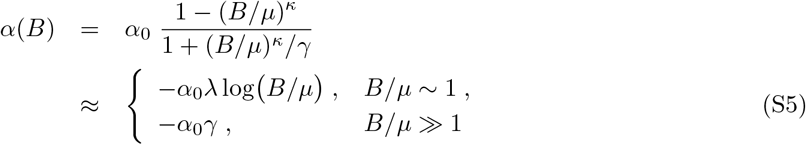

with *λ* = *κγ/*(1 + *γ*). Both approximations are illustrated in Fig. S1. Previous work has mainly considered antibiotic concentrations where the linear regime is negligibly small and the curve can be approximated by a step function [19]. We are here interested also in the tug-of-war dynamics that occurs around the intermediate regime of antibiotic concentrations approximately linear. When deriving results below, we will compare these two regimes. Specifically, section A.1 deals with the constant approximation for large enough antibiotic concentrations, while section A.2 treats the logarithmic approximation close to *B*_0_ ≈ *µ*.

### A.1 Solutions to different decay mechanisms with constant death rate

At first, we treat the regime with a constant death-rate, *α*(*B*) ≈ −*α*_0_*γ*. In the ensuing calculation, we rescale time,

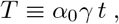

such that bacterial death rate is 1 in these dimensionless units. With this rescaling, the population dynamics is simply *∂*_*T*_ *N* = −*N*. We immediately find the solution for the microbial population as *N* (*T*) = *N*_0_ exp(−*T*). This gives the time *T*_1_ = log *N*_0_ for survival in antibiotics: at this time, only one cell would remain, (*N* (*T*_1_) = 1, if antibiotics are still present, and the population is essentially extinct. The exponential decay of the population size allows to solve the dynamics of the antibiotic concentration in other cases as well. In order to obtain these solutions for the antibiotic dynamics, we introduce its logarithmic concentration,

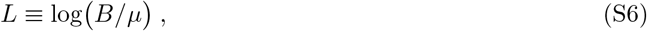

which simplifies the dynamics, as in all kinetic cases defined above the dynamics of antibiotics itself is linear in *B*, see Eqs. (S3). We multiply each equation with 1*/B*, and use the identity 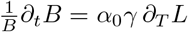. The change of time units from *t* to *T* for the derivation introduces the coefficient *α*_0_*γ*. Using the logarithmic antibiotic concentration *L*, and inserting the solution *N* (*T*) = *N*_0_ exp(−*T*) for the population size, we can integrate all trajectories *L*(*T*), and obtain

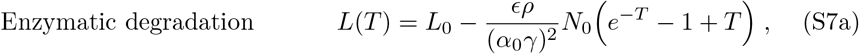

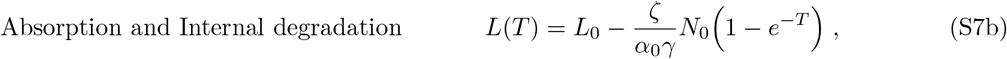

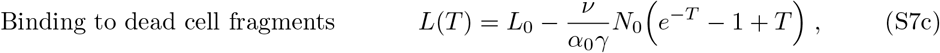

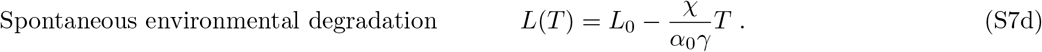

In these solutions, we observe that every parameters occurs in a single coefficient, and thus can be combined into an effective and dimensionless parameter. We define the timescale separation *τ* for each of the different degradation mechanics as,

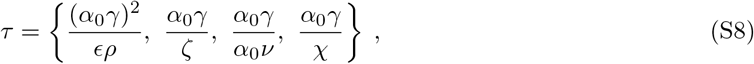

in the same order as above. Independent of the mechanism, *τ* is always the ration of the rate for bacterial death over the rate of antibiotic degradation. In the case of enzymatic degradation, which consists of two steps with enzyme production and then degradation, the death rate is squared to make *τ* dimensionless.

Now checking for population survival, we evaluate the antibiotic concentration *L*(*T*_1_) at time *T*_1_ = log *N*_0_. If *L*(*T*_1_) < 0 (which translates to *B*(*T*_1_) < *µ*), then the antibiotic concentration has been reduced far enough, that the population reached already positive values for its growth rate *α*(*B*), and started growing before it reached a single cell. Conversely, if still *L*(*T*_1_) *>* 0, then the population would decay more and go extinct. Inserting *T*_1_ = log *N*_0_ into each of the solutions in Eq. (S7), evaluating the condition *L*(*T*_1_) < 0, and finally reverting back to *B*_0_ from *L*_0_ yields the expressions

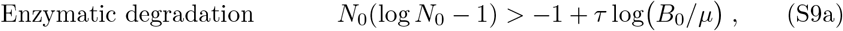

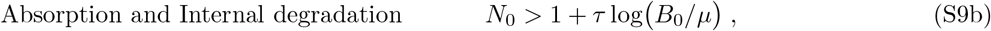

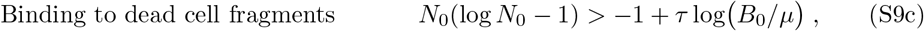

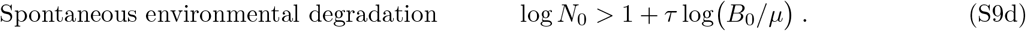

to antibiotic concentration *B*_0_ instead of *L*_0_. With the exception of spontaneous environmental degradation, we neglect the logarithmic population size dependence, and arrive at the universal expression for the MSPS curve,

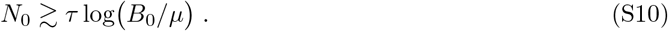

### A.2 Solutions by linear approximation close to MIC

Now we focus on the dynamics of competition when the initial antibiotic concentrations is intermediate, close to *B*_0_*/µ* ≈ 1. In this regime, antibiotic-dependent growth rate is approximated with the linear expansion in (S5):

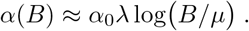

We utilize again a rescaling of time and the logarithmic antibiotic concentration

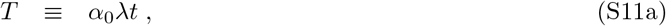

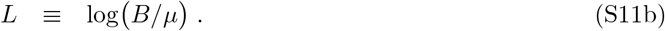

Then, the equation for population dynamics takes the form

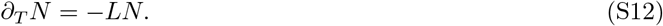

For each of the four reactions specified above, we define a dimensionless parameter *τ*, given by (in order of the reactions)

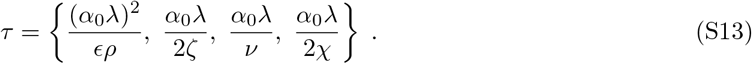

These parameters are the again ratios of timescales between population death and antibiotic degradation. Due to the coupling of antibiotic concentration and population size dynamics in *∂*_*T*_ *N* = −*LN*, solving these dynamics is more involved as before. Thus, we treat each mechanism is a separate section in the following.

#### A.2.1 Enzymatic degradation

For enzymatic degradation, in the newly defined variables the equations are

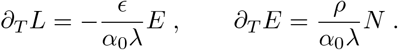

Differentiating the first equation again with respect to *T*, then using the second equation for *∂*_*T*_ *E*, we find two coupled equations for population size and antibiotics

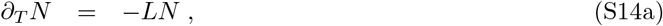

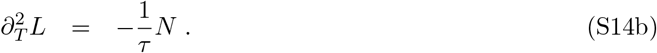

We cannot state explicit closed form solutions for this case. Nevertheless, we can assume a constant exponential decay of the bacterial population, which is most of the time is close to the exact solution (checked numerically). Only close to the turning point (where bacteria start to grow again due to degraded antibiotics), we expect significant deviations from this simple exponential decay.

To this end, we use *N* (*T*) ≈ *N*_0_ exp (−*L*_0_*T*), which can be integrated twice to obtain the trajectory *L*(*T*). Initial conditions for the latter are (*∂*_*T*_ *L*)(0) = 0 (no enzyme present at the beginning) and *L*(0) = *L*_0_. The solutions are

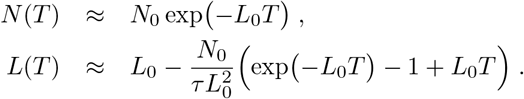

Estimating the time *T*_1_ = (log *N*_0_)*/L*_0_ from inverting *N* (*T*_1_) = 1, we compute the condition for survival of the bacterial population again from *L*(*T*_1_) < 0. Algebraic rearrangements of this inequality lead to

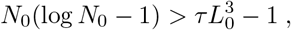

which in turn can be expressed in the Lambert-W function 𝒲 (also called product-logarithm),

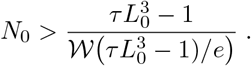

For large values of 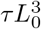 (note that *τ* ∼ 𝒪(10^6^), cf. Fig. 5), we have *W*(*X*) *∼* log *X*. This logarithmic correction is usually small, and we neglect it to arrive at the MSPS curve,

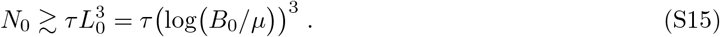

However, in contrast to before we find an exponent 3 for the (logarithmic) antibiotic concentration. In terms of our experiments, this would introduce a sharper bend on the plate close to the transition to the vertical part. For larger initial concentrations *B*_0_, however, the MSPS curve transitions into the solution with constant death rate treated before.

#### A.2.2 Absorption and internal degradation

The dynamics for absorption and internal degradation can be solved analytically. This mechanism assumes a first order mass-action absorption reaction of antibiotics to cells, and then immediate degradation of antibiotics inside the cell (or at least very fast compared to absorption). The system of differential equations is given by

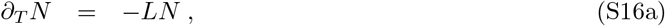

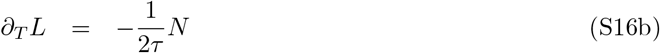

where *τ* is the second definition in (S13). Upon taking the ratio of both of these equations, we find 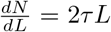. This auxiliary differential equation can be integrated to arrive at *N* (*L*) = *τ L*^2^ + *a* with *a* an integration constant. Consequently, (S16b) becomes *∂*_*T*_ *L* = −*L*^2^*/*2 − *a/*(2*τ*). Although non-linear, this differential equation has a solution in trigonometric functions, and we have after inserting *L*(*T*) into *N* (*L*) the two solutions

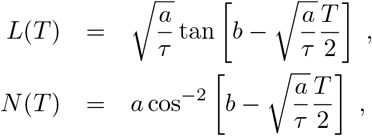

where *b* is the second integration constant. Using the initial conditions *N* (0) = *N*_0_ and *L*(0) = *L*_0_, we find

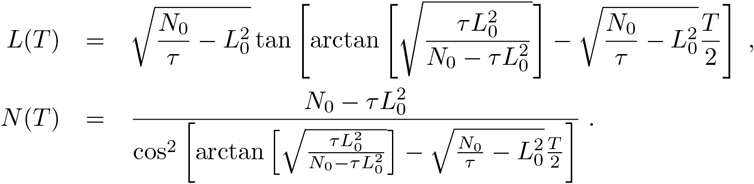

Although these expressions look complicated, we can utilize the special trigonometric functions to proceed. The arguments of tan and cos^−2^ in these solutions are identical, and we know that the zeros of tan correspond to minima of cos^−2^ with value 1 ((tan[*X*] = 0) ⇔ (cos^−2^[*X*] = 1), *∀X*). Thus, we know that the time *T*_*L*_, defined as *L*(*T*_*L*_) = 0 leads to 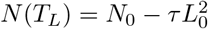, which we have to check if it is bigger or smaller than one. Consequently, we obtain the MSPS curve

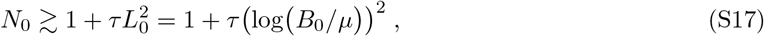

from rearranging this condition.

#### A.2.3 Irreversible binding to cell fragments

Another degradation mechanism is irreversible binding of antibiotics to dead cell fragments and their consequent inactivation. The dynamic equations in this case are

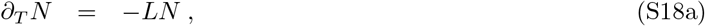

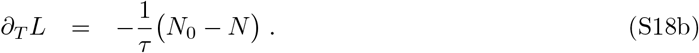

We approximate an exponential decay as solution to the bacterial population size, and then integrate the second equation for the (logarithmic) antibiotic concentration. This yields

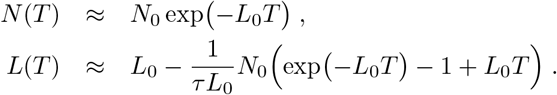

Using these solutions, we again check *L*(*T*_1_) < 0, which translates into the condition on the minimal population size

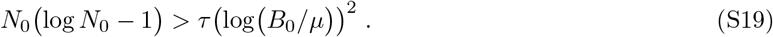

#### A.2.4 Spontaneous environmental degradation

As the simplest of all cases, we treat spontaneous environmental degradation. The set of two equations is given by

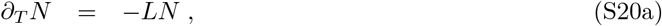

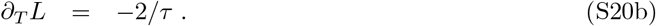

This system can be solved almost trivially: We immediately find *L*(*T*) = *L*_0_ − 2*T/τ*, which we can insert into the dynamics of *N* to find *N* (*T*) = *N*_0_ exp *T*^*2*^ */τ L*−*T)*. This solution for the population size has a minimum at time *T*_min_ = *τ L*_0_. Thus, the population will survive if 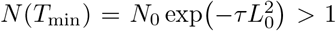. Reverting back to original variables yields the condition

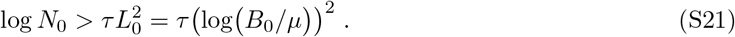

Here, we again find a logarithmic dependence on initial population size *N*_0_, which deviates from the MSPS curve.

Overall, we have seen in this section, that for initial antibiotic concentrations close to *µ*, the exponent in the MSPS curve increases. This leads to a sharper bend away from the vertical part (see Fig. 3B). However, on plates which increase the initial antibiotic concentrations fast enough along one axis, we soon arrive at a dynamic where the constant approximation treated in section A.1 is more appropriate.

## B Privatization effects of an extracellular enzyme

The main text only contained an overview of the steps involved in deriving a model that includes dynamical privatization. Here, we present a more detailed account of these computations.

Our original model included bacterial growth, together with production of an enzyme (eg. *β*-lactamase) that reduces antibiotics. Now, we include additional ODEs explicitly describing the time-evolution inside and outside of cells,

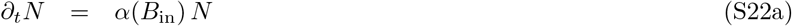

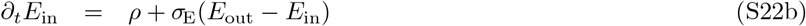

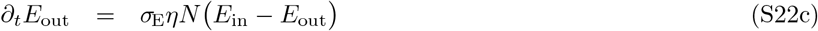

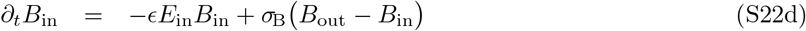

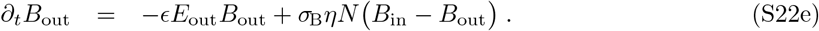

Enzyme *E* and antibiotics *B* are transported between the inside and the outside with rates *σ*_E_ and *σ*_B_, respectively. The volume-separation factor *η* indicates how much slower the concentrations outside cells change, compared to the rather fast dynamics within. This factor occurs, as the volume of the culture medium *V*_medium_ is usually much larger as the volume of all cells *V*_cells_, and we have to account for the fact that a single molecule passing through the membrane affects concentrations in different ways for internal and external concentrations. Thus, we need to couple the transport processes via a factor *V*_cells_*/V*_medium_ = *ηN*, where *η* is the ratio of volumes for a single cell. As long as *η* ≪ 1, we can assume that adjustment of inner concentrations (*E*_in_ and *B*_in_) is essentially instantaneous compared to the dynamics outside cells. Explicitly, for our experiments we have *V*_medium_ = 2 · 10^−4^*mL* and *V*_cell_ ≈ 10^−15^*mL*, such that *η* ≈ 5 · 10^−12^. Hence, we can make an approximation that these internal concentrations are stationary, and set both *∂*_*t*_*E*_in_ ≈ 0 and *∂*_*t*_*B*_in_ ≈ 0. This allows to algebraically rearrange Eqs. (S22b) and (S22d) to express inner concentrations as functions of the (slower varying) outer concentrations,

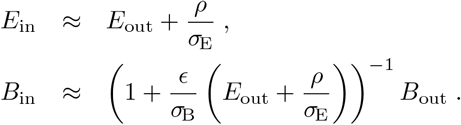

These relations indicate that the effective enzyme concentration inside cells is higher by an additive term *ρ/σ*_E_ compared to outside cells. Moreover, assuming the outside enzyme concentration remains small relative to inside, *E*_out_ *« ρ/σ*_E_ the effective antibiotic concentration is reduced by a factor 1 + *cρ/σ*_E_*σ*_B_.

Inserting these two expressions in the outside dynamics leads to

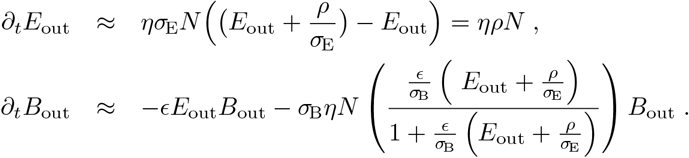

In the first line, we recover the expected dynamics of enzyme production, *∂*_*t*_*E* = *ρηN*, where the original production rate *ρ* needs to be adjusted to also incorporate the volume-separation factor *η*. The dynamics of antibiotics can be rewritten as *∂*_*t*_*B* ≈ −ϵ(*E*_out_ + *σ*_B_*ηN*)*B*, which now also exhibits a second term similar to the case of ‘absorption and internal degradation’, described above in Eq. (S3b) and section A.2.2. There, we effectively had 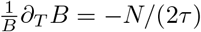. However, we expect that this new additional term proportional to *σ*_B_*ηN* only affects the dynamics late into the experiment, when many cells are already present. This fact is corroborated when inserting extracted values for parameters from our experiments, where the first coefficient is 1*/τ ∼ O*(10^−6^) from which we can deduce that the second coefficient appearing now in the antibiotic degradation is *O*(10^−9^). All these considerations change the timescale separation *τ* for the explicit treatment of internal concentrations to

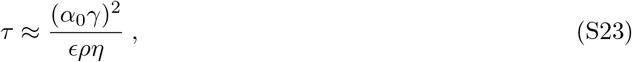

which contains an additional factor *η*. Note, however, that this does not change the value of *τ* itself, but rather which values we have to attribute to the microscopic parameters *ρ* (and maybe *ϵ*) when conducting simulations that are supposed to match either experiments or simulations for the dynamics without these explicit inner concentrations.

So far, we always measured antibiotic concentrations *B* in units of multiples of *µ*. The dynamics of *B* itself, see Eqs. (S22d) and (S22e), is invariant under multiplication with a scale factor, that can be chosen arbitrarily (and we used *µ* to fulfill this role). The only point in our model, where we need proper units for the antibiotic concentrations, is to quantify the effects on growth rate, defined via Eq. (2), where the ratio *B/µ* enters. Moreover, in the derivation of privatization effects above, we found that the cell effectively sees a slightly reduced concentration of antibiotics, *B*_in_, as can be measured in media, *B*_out_. Thus, we can write

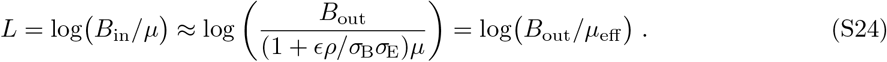

This allows to state the overall dynamics in the outer concentration *B*_out_ as we did in Eq. (S24), we can now relate the effective single cell MIC *µ*_eff_ to the underlying dynamical parameters. In the main text we introduced the privatization parameter Φ = (*ϵρ*)*/*(*σ*_B_*σ*_E_) that occurs here. Writing Φ in terms of the timescale separation *τ*) in Eq. (S23), we obtain a parameter *φ* characterizing the privatization effect,

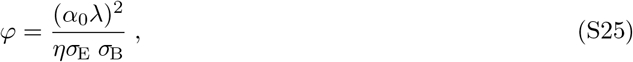

such that Φ = *φ/τ*.

Specifically, we find that the antibiotic concentration where population growth turns into death in resistant populations is given by

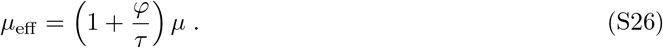

This relation (S26) is linear in 1*/τ*, which is the one parameter characterizing an allele. In general, we expect *φ* to be a constant for both our sets of experiments.

Susceptible cells should correspond to a limit of *τ →* 0, as then either production is negligible (*ρ →* 0) or the (non-existent) enzyme is not effective (*ϵ →* 0). As a sanity check, extrapolating the scMIC values to *τ* = 0 should give the *µ* value measured in kill curve experiments.

### C Quantification of bacterial growth with antibiotic dosage

Methods used to estimate the impact of antibiotic levels on the growth of bacterial cultures – referred to as “kill curves” – were adapted from [23]. Briefly, *Escherichia coli* MG1655 galK::SYFP2-FRT was cultured overnight in M9 minimal media (supplemented with 0.4% glucose, 0.2% Casamino acids, 1 *µg/mL* thiamine, 2 *µg/mL* uracil and 50 *µM* IPTG). Stationary phase cultures were diluted 1:1000 into minimal media and incubated shaking, meanwhile dilution series were made of cefotaxime in minimal media, 140 *µL* of each dilution was aliquoted into a row of a 96-well plate. All media for subsequent culturing was then pre-warmed to aid continuous growth of bacteria. After 120 minutes, the concentration of cells was measured by flow cytometry, and diluted in minimal media to approximately 2 · 10^6^ cells*/mL*, and 140 *µL* of cells were added to each well containing cefotaxime. A 20 *µL* sample was immediately taken, and the plate was incubated with orbital shaking for 60minutes at 37°C with further 20 *µL* samples taken every 10 minutes. For each sample a dilution series of 10^−1^ to 10^−3^ was made and 100 *µL* of each dilution was plated on minimal media solidified with agar. Plated cell solution were spread with 12-14 3 *mm* glass beads. Plates were incubated for *∼* 24 *h* and colony forming units were counted from appropriate dilutions.

## D Experimentally determining the MSPS curve

### D.1 Detailed protocol to perform 96-well microtitre plate assay

The proposed assay exposes different inoculum sizes of bacteria to different concentrations of antibiotics until the boundary between wells with surviving and extinct populations is evident. The population size is varied across the eight rows (A-H) of a 96-well plate while antibiotic concentrations varies across the twelve columns of the plate. Care should be taken when choosing the antibiotic dilution range to fully capture the MSPS curve. If the standard MIC is known, the dilutions series may be chosen so that the MIC is the 8th or 9th dilution of this dilution series.

1. Grow strain of interest overnight in a suitable growth medium.
2. Prepare an antibiotic solution in the chosen growth medium that is four times the highest concentration that will be tested.
3. Fill all wells in a sterile 96-well plate with 100 *µL* growth medium.
4. Dispense 100 *µL* of the antibiotic solution with a concentration 4X of the intended final concentration in column one of the microtitre plate. Using a multi-channel pipette, mix the antibiotics and transfer 100 *µL* from column one to column two. Mix again and repeat this procedure down to column 12. Discard 100 *µ*l solution from column 12.
5. Prepare a serial dilution of the bacterial overnight culture in eight appropriate tubes. The highest concentration should be twice the desired highest inoculum. The highest inoculum size in the validation experiments was approximately 1.25 · 10^6^ CFU*/ml* or 2.5 · 10^5^ CFU*/*well.
6. Dispense 100 *µL* of this serial dilution series across the rows of 96-well plate.
7. Incubate the plates at 37°C for 24h or until satisfactory growth is obtained.
8. Growth can be visually assessed or by spectrophotometric reading (OD600).

The validation experiments described in this study were performed using the above-mentioned protocol. For the set of experiments using enzyme variants with a different catalytic activity, TEM-1 and three alleles (G238S, E104K-G238S, E104K-M182T-G238S) were amplified from previously described plasmid constructs [30], and introduced into chromosomal *galK* of *Escherichia coli* MG1655 by using the ‘Quick and Easy E. coli Gene Deletion Kit’ (Gene Bridges). Mutants were selected by selection for ampicillin resistance and the introduction of *β*-lactamase genes were confirmed by PCR and sanger sequencing. The MSPS assay was performed with the same minimal media used for kill-curves. For the set of experiments assessing the effect of expression, TEM-1 alleles G238S and E104K-M182T-G238S were subcloned into pBAD322T behind an arabinose-ineducable promoter and transformed in *E. coli* BW27783 (CGSC#12119) which carries a deletion for the arabinose metabolizing genes [31, 32]. Here, the MSPS curve was performed in standard LB medium supplemented with 0.1% (High Expression), 0.003125% (Medium Expression) 0.00078125% (Low Expression), and 0% (No/Leaky Expression) L-Arabinose where appropriate. For both sets of experiments, growth was measured after 24h using a Victor3 plate reader (Perkin-Elmer).

### D.2 Parameter estimation on plates

Estimating the two parameters *τ* and *µ*_eff_ from the OD of one plate involves multiple steps. First, we interpolate the 12 *×* 8 grid of the 96-well plate onto a much finer grid with a Gaussian Process Regression (GPR). Using this finer grid, we estimate a threshold between growth and no-growth by applying Otsu’s method to find a threshold value that separates the modes in a bi-modal distribution of OD values. We compute the contour line in the fine grid at this threshold value to obtain points on a curve close to the growth/no-growth threshold. In the last step we fit the the predicted MSPS curve, *N*_0_ ≈ *τ* log (*B/µ*_eff_), to this contour line, which yields values for *τ* and *µ*_eff_. The main concepts and equations of all steps are described in more detail below.

The implementation in Python can be found on https://github.com/lukasgeyrhofer/antibiotics.

#### D.2.1 Otsu’s Method for threshold detection

The next step in extracting the parameters from plates is to estimate the threshold between growth and no-growth in an automated way. We use a slightly modified version of Otsu’s method [33] for finding this threshold. Originally, this method has been proposed for binarizing images into black and white by separating the all observed values into two classes. It works by minimizing variance within a class, or equivalently, by maximizing variance between the two classes.

In order to compute the threshold, we first sort the OD values from all replicate plates into a vector **a** of length *M · R* (with *M* = 96 the number of OD values and *R* the number of replicates),

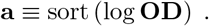

Here, we use logarithmic values, as usually only the magnitude of the growth and no-growth OD values is important, which allows to get a clearer threshold value. Then, one can show that Otsu’s method can be reduced to finding the index *m* that maximises the expression,

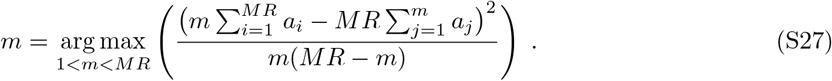

Originally, Otsu’s method used equally distant bins and counts the number of values within these bins. However, in our modified version, Eq. (S27), we use ‘bins’ with only one observation, and space them according to the actual measurements.

The threshold between growth and no-growth is then defined as 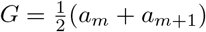. All OD values above this threshold *G* are considered growing, while all values below are considered no-growth.

#### D.2.2 Parameter estimation

With the set of points for the threshold between growth and no-growth, obtained in the previous step, we first transform both axes to their logarithmic scales, which gives a set of the initial conditions (log(*N* − 1))_*i*_, (log *B*)_*i*_ for each of the wells. Then we fit the (non-linear) functional form

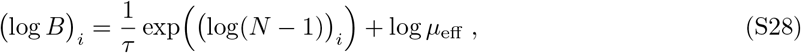

to first obtain values for (1*/τ*) and intercept (log *µ*_eff_) using Python’s *scipy*.*optimize*.*curve fit*. Then, using the Python module *uncertainties*, we use the covariance matrix between these two fitted parameters to compute the actual values *τ* and *µ*_eff_, in addition to their standard deviations, which are shown in Fig. 5. The advantage of using logarithmic values for both axes of initial population size and antibiotic concentration is that distances are weighted according to distances on the plate. Evaluating the exp(log *N*_*i*_) term to obtain the direct linear dependency on *N* would emphasize large initial populations too much in the fitting procedure.

## E Supplemental Figures

**Figure S2:**
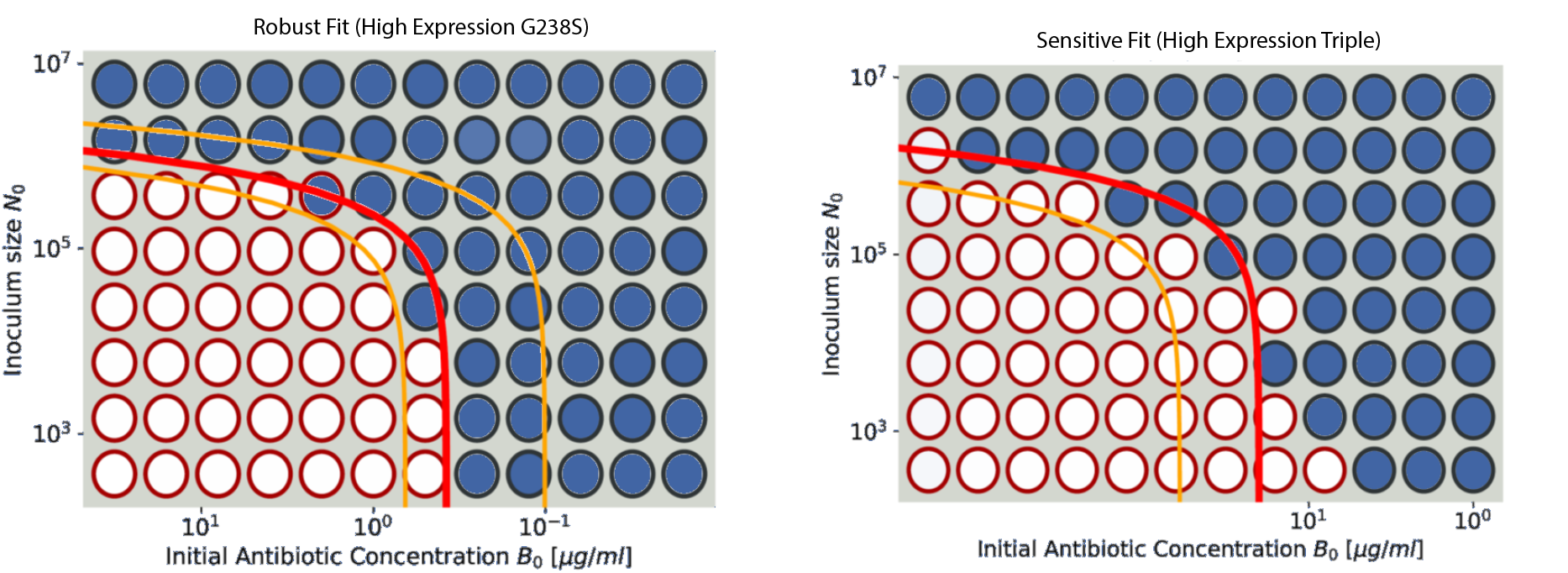
Individual fits from microtitre plate experiments showing a robust and sensitive fit of the MSPS curve.

